# Rhamnose biosynthesis is not impaired by the deletion of putative *rfbC* genes*, slr0985* and *slr1933*, in *Synechocystis* sp. PCC 6803

**DOI:** 10.1101/2025.03.27.645739

**Authors:** João Pissarra, Marina Santos, Sara B. Pereira, Catarina C. Pacheco, Filipe Pinto, Sónia S. Ferreira, Ricardo Monteiro, Cláudia Nunes, Manuel A. Coimbra, Didier Cabanes, Rita Mota, Paula Tamagnini

## Abstract

Cyanobacterial extracellular polymeric substances (EPS), mainly composed by heteropolysaccharides, can be attached to the cell wall (CPS) or released to the environment (RPS). These polymers have an unusually highly diversified monosaccharidic composition, making them attractive for biotechnological/biomedical applications. However, their production is still poorly understood hindering their optimisation for industrial needs. This work aimed at better understanding the biosynthesis of the 6-deoxysugars fucose and rhamnose in the model cyanobacterium *Synechocystis* sp. PCC 6803. To that end, genes encoding proteins putatively involved in the biosynthesis of GDP-L-fucose [*sll1213* (*fucS*)] and dTDP-L-rhamnose [*slr0985* (*rfbC1*) and *slr1933* (*rfbC2*)] were deleted. As previously observed, Δ*fucS* had significant growth impairment and its RPS did not contain any fucose or rhamnose. Here, we also showed that both deoxyhexoses’ pathways are completely impaired in Δ*fucS*. In contrast, both Δ*rfbC1* and Δ*rfbC1*Δ*rfbC2* although producing significantly less RPS and more CPS than the wild type, did not show major differences regarding the RPS monosaccharidic composition. These results strongly suggest that their gene products are not essential for rhamnose biosynthesis. Transcriptional analysis revealed that one of the *gmd* genes (*slr1072*), putatively encoding a GDP-mannose 4,6-dehydratase, was upregulated in all the knockout strains, and that the three EPS-related genes in the same operon as *rfbC1* (*slr0982*, *slr0983* and *slr1610*) were upregulated in both Δ*rfbC* strains. Altogether, our results reveal that rhamnose biosynthesis in *Synechocystis* depends on FucS but not on the putative RfbC enzymes, underlining the need to further elucidate the mechanisms involved in the biosynthesis of this deoxyhexose.

**IMPORTANCE:** This study contributes to the overall knowledge of deoxyhexoses’ biosynthesis in *Synechocystis* sp. PCC 6803. Here we demonstrated that the Δ*fucS* strain not only produces EPS without fucose and rhamnose but that both pathways are completely impaired. Furthermore, we also showed that the deletion of both putative *rfbC* genes do not affect rhamnose biosynthesis, despite having an impact on carbohydrates production/export, shifting RPS to CPS production. Altogether, our results suggest that the *rfbC* genes are not correctly annotated and highlight the intricacies and/or potential crosstalk between the two deoxyhexoses pathways, yet to be completely unravelled in *Synechocystis*. The understanding of cyanobacterial EPS assembly and export is crucial for the optimisation of their production and tailoring for industrial/commercial applications.

## INTRODUCTION

Many cyanobacteria can produce extracellular polymeric substances (EPS) that are mainly composed by heteropolysaccharides. EPS can remain attached to the cell surface as capsular polysaccharides (CPS) or be released to the extracellular medium as released polysaccharides (RPS) (Rossi and De Philippis, 2016). Several biological functions can be attributed to EPS, such as protection against several external factors, nutrient sequestration, water retention, carbon reservoirs, intercellular interactions, formation of biofilms, motility, and cell adhesion (Kehr and Dittmann, 2015; Li et al., 2021; Rossi and De Philippis, 2015). The interest on cyanobacterial EPS for biotechnological applications – mostly in high value markets like the cosmetic and pharma industries – is increasing as an alternative to synthetic polymers, as well as due to their potential bioactivities as antiviral, antibacterial, antifungal, antioxidant, anti-inflammatory, antitumor, and anticoagulant (Flamm and Blaschek, 2014; Flores et al., 2019; Morales-Jiménez et al., 2020; Mota et al., 2023; Pereira et al., 2019; Reichert et al., 2017). However, cyanobacterial polymers must compete with polymers already established in the market mainly due to scalability and productivity issues, and/or processing costs (Agarwal et al., 2022; Gomes Gradíssimo et al., 2020; Laroche, 2022; Pierre et al., 2019). On top of that, the potential of the cyanobacterial polymers will only be fully exploited when their biosynthetic pathways are better understood, paving the way to tailoring them to industrial needs (Mota et al., 2021; Pereira et al., 2019).

In contrast to other bacterial polymers, cyanobacterial EPS can be composed by up to 13 different monosaccharides, including hexoses (glucose, mannose, galactose, and fructose), deoxyhexoses (fucose and rhamnose), pentoses (arabinose, ribose, and xylose), acidic (glucuronic and galacturonic acid) and amino sugars (glucosamine and galactosamine), as well as sulphate, methyl and acetyl groups, and even peptides and several other non-carbohydrate constituents, resulting in complex and highly variable structures (Delattre et al., 2016; Flores et al., 2019; Kehr and Dittmann, 2015; Madsen et al., 2023; Mota et al., 2021, 2020; Pereira et al., 2009, 2019; Tiwari et al., 2020). In several organisms, the 6-deoxysugars play an important role in maintaining the integrity of the cell wall and capsule, and are incorporated into glycoproteins and several glycosylated metabolites such as antibiotics (Bischer et al., 2020; Carvalho et al., 2018; Mäki and Renkonen, 2004; Mohamed et al., 2005; Reiter, 2002). These sugars may also confer bioactive properties to EPS, since antioxidant, antibacterial and biofilm-inhibiting properties have been associated to fucose-rich bacterial EPS (Baptista et al., 2023; Ramachandran et al., 2020). The polysaccharide fucoidan isolated from the cell wall of brown algae has also been linked to antitumor, antioxidant, antiviral, and anti-inflammatory activities (Wang et al., 2019). Bacterium-, plant-, and synthetically-derived glycosides containing rhamnose have been linked to antitumoral, antibacterial and even anti-inflammatory activities (Chen et al., 2018; Lee et al., 2014; Mei et al., 2019; Sylla et al., 2019), while bacterial EPS containing rhamnose have been found to have antioxidant and biofilm-inhibiting properties (Xu et al., 2020). In cyanobacterial EPS, fucose and rhamnose are mostly assumed to confer hydrophobic properties that can contribute to cell adhesion and polymers emulsifying properties (Pereira et al., 2019; Rossi and De Philippis, 2015). Deoxyhexoses are often only biologically available in activated forms with nucleotide groups, two of the most common being guanosine diphosphate-L-fucose (GDP-Fuc) and deoxythymidine diphosphate-L-rhamnose (dTDP-Rha) (Li et al., 2022; Mäki and Renkonen, 2004). Despite being extensively characterised in several organisms, most of what is known regarding the activation and conversion of sugar nucleotides in cyanobacteria is inferred (Han et al., 2017; Mills et al., 2020; Santos-Merino et al., 2023). The *de novo* biosynthetic pathway of GDP-Fuc is highly conserved among bacteria, plants, fungi and animals, and entails a three-step reaction catalysed by two enzymes, GDP-mannose 4,6-dehydratase (Gmd) and GDP-L-fucose synthase (FucS, also known as WcaG or GMER). In this reaction, GDP-D-mannose is oxidised and dehydrated into GDP-4-keto-6-deoxy-D-mannose that is subsequently epimerised into GDP-4-keto-6-deoxy-L-galactose and finally reduced into GDP-Fuc (Mäki and Renkonen, 2004; Niittymäki et al., 2006; Ren et al., 2010). The genes encoding Gmd and FucS are generally clustered and are nearly ubiquitous (Mäki and Renkonen, 2004), with cyanobacteria being no exception. Several putative orthologues encoding these two proteins have been described in many cyanobacterial strains like *Synechocystis* sp. PCC 6803, *Synechococcus* sp. PCC 7002, *Thermosynechococcus elongatus* BP-1, *Anabaena* sp. PCC 7120 and *Nostoc punctiforme* ATCC 29133 (Graham and Bryant, 2009; Mochimaru et al., 2008; Mohamed et al., 2005). According to the biosynthetic pathways described for *Synechocystis* sp. PCC 6803 in the Kyoto Encyclopedia of Genes and Genomes (KEGG; Kanehisa, 2019; Kanehisa et al., 2023; Kanehisa and Goto, 2000) and CyanoCyc (Moore et al., 2024) databases, and in a review by Mills et al. (2020), there are two candidate genes, *sll1212* and *slr1072*, encoding the GDP-mannose 4,6-dehydratase (Gmd), while FucS is encoded by *sll1213*. It has also been shown that *fucS* deletion leads to a distinct phenotype, including impaired growth, cell aggregation, lack of S-layer, altered organisation of the thylakoid membranes, EPS without fucose and rhamnose, lipopolysaccharides with truncated O-antigen portions, abnormal vesicle size, and hypervesiculation (Fisher et al., 2013; Matinha-Cardoso et al., 2022; Mohamed et al., 2005). The biosynthetic pathway of dTDP-Rha is highly conserved and consists of four steps, each catalysed by a different enzyme, namely glucose-1-phosphate thymidylyltransferase (RfbA or RmlA), dTDP-glucose 4,6-dehydratase (RfbB or RmlB), dTDP-4-dehydrorhamnose 3,5-epimerase (RfbC or RmlC), and dTDP-6-deoxy-L-mannose-dehydrogenase (RfbD or RmlD). Firstly, the nucleotide thymidylmonophosphate is transferred onto the substrate glucose-1-phosphate. Then, the D-glucose residue is oxidised and dehydrated, leading to the formation of dTDP-4-keto-6-deoxy-D-glucose, followed by a double epimerisation resulting in dTDP-4-keto-6-deoxy-L-mannose. Finally, the C4 keto group is reduced, forming dTDP-Rha (Giraud and Naismith, 2000; Li et al., 2022; Mäki and Renkonen, 2004). The genes encoding RfbA, RfbB, RfbC and RfbD are often scattered and while many putative orthologues have been identified in cyanobacterial strains, such as *Synechocystis* sp. PCC 6803, *Prochlorococcus* spp. and *Anabaena* sp. PCC 7120 (Huang et al., 2005; Mills et al., 2020; Sun and Luo, 2018), others seem to lack the pathway for the biosynthesis of dTDP-Rha (Burgsdorf et al., 2015; Mills et al., 2022). In *Synechocystis* sp. PCC 6803, RfbA is likely encoded by *sll0207*, RfbB by *slr0836*, and RfbD by *sll1395*. However, there are two *rfbC* candidates, *slr0985* and *slr1933* (Kanehisa, 2019; Kanehisa et al., 2023; Kanehisa and Goto, 2000; Mills et al., 2020; Moore et al., 2024).

In this study, we aimed at better understanding the pathways involved in the biosynthesis of the deoxyhexoses GDP-Fuc and dTDP-Rha in *Synechocystis* sp. PCC 6803, contributing to unveil the intricate mechanisms of cyanobacterial EPS assembly and export. For this purpose, we generated Δ*slr0985* (Δ*rfbC1*) and Δ*slr0985*Δ*slr1933* (Δ*rfbC1*Δ*rfbC2*) *Synechocystis* knockout strains and characterised them in relation to the previously generated Δ*sll1213* (Δ*fucS*, Matinha-Cardoso et al., 2022) and the wild type strain. Characterisation included growth analyses, EPS production and monosaccharidic composition, ultrastructure of the cell envelope, as well as analysis of the transcript levels of target genes.

## RESULTS

### Genomic context of *sll1213* (*fucS*), *slr0985* (*rfbC1*) and *slr1933* (*rfbC2*) in *Synechocystis* sp. PCC 6803 and deletion strains’ generation

This work aimed at better understanding the biosynthesis of the 6-deoxysugars fucose and rhamnose in the model cyanobacterium *Synechocystis* sp. PCC 6803 (hereafter *Synechocystis*), and the impact of the absence of putative key enzymes, namely the GDP-L-fucose synthase (FucS) and the dTDP-4-dehydrorhamnose 3,5-epimerase (RfbC), on EPS production (schematic representation of the putative pathways and locus of the respective genes are depicted in Fig. 1). For this purpose, three strains were used: Δ*sll1213* (Δ*fucS*, Matinha-Cardoso et al., 2022), Δ*slr0985* (Δ*rfbC1*) and Δ*slr0985*Δ*slr1933* (Δ*rfbC1*Δ*rfbC2*). The last two strains were generated within this study and their segregation was confirmed by PCR and Southern blot (Fig. S1).

**Fig. 1.**
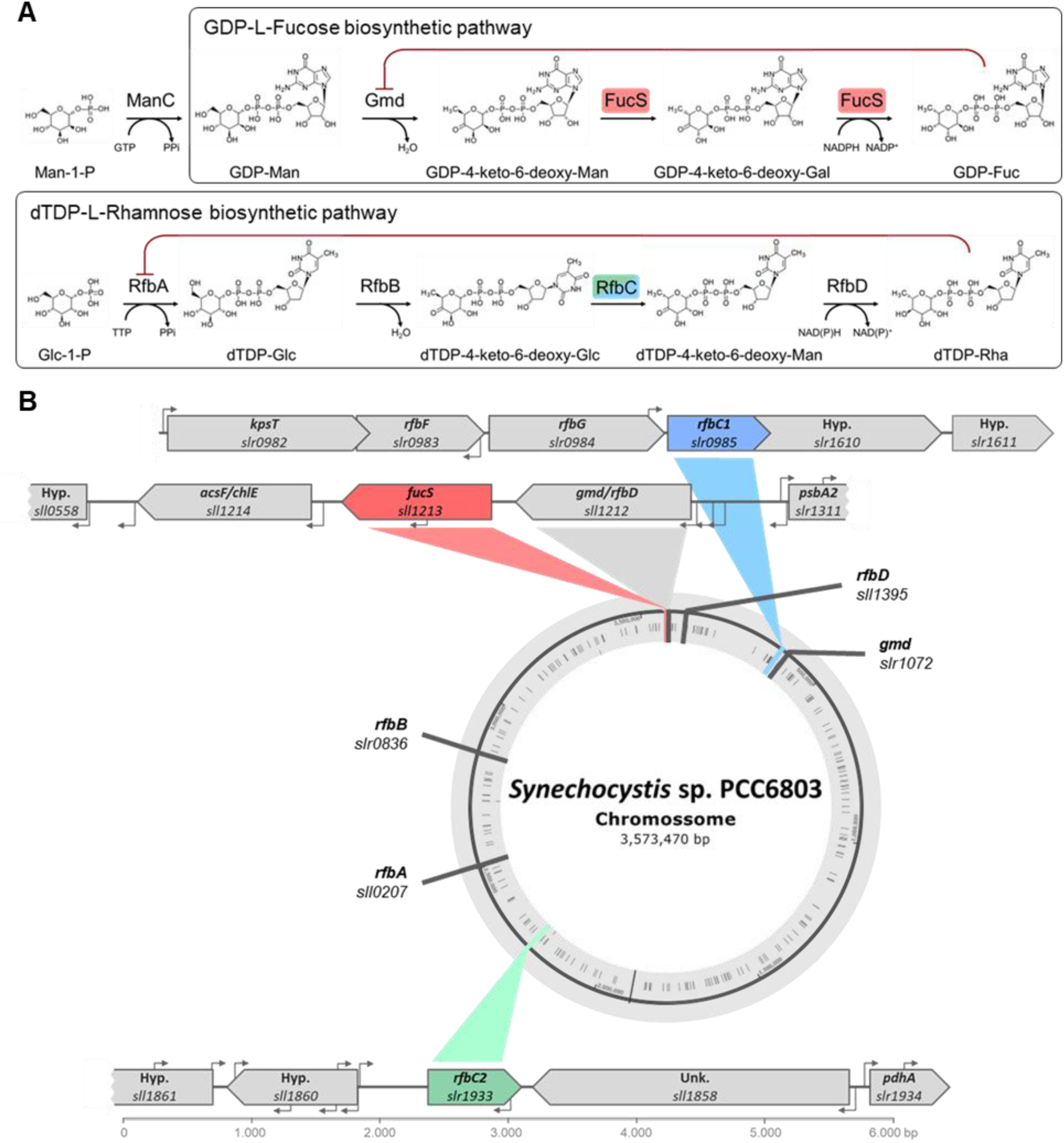
Biosynthetic pathways and genomic context of genes involved in the synthesis of fucose and rhamnose in *Synechocystis* sp. PCC 6803. (A) Schematic representation of the putative biosynthetic pathways of guanosine diphosphate-L-fucose (GDP-Fuc) and deoxythymidine diphosphate-L-rhamnose (dTDP-Rha). The red line indicates feed-back inhibition. (B) Locus of the genes encoding enzymes putatively involved in the biosynthetic pathways of GDP-Fuc and dTDP-Rha, with the genes knockout in this study highlighted in red, blue and green. Both the locus tag and gene name/symbol(s) are provided when available. Unk./Hyp. - genes encoding unknown or hypothetical proteins.

### Growth, S-layer and carbohydrates production

To understand how the deletion(s) of the selected genes affect *Synechocystis* features and carbohydrates production, the strains were grown in liquid BG11 medium. Remarkably, the Δ*fucS* strain displayed an accentuated clumping phenotype at low optical densities (OD_730_ < 1), while the Δ*rfbC* strains did not exhibit this obvious phenotype (Fig. 2A). In addition, all strains showed faster sedimentation index compared with the wild type (Fig. 2B). However, while 85% of the Δ*fucS* cells and 81% of the Δ*rfbC1*Δ*rfbC2* cells have sedimented after 24 h, only 35% of the Δ*rfbC1* cells had sedimented at this time point (Fig. 2B).

**Fig. 2.**
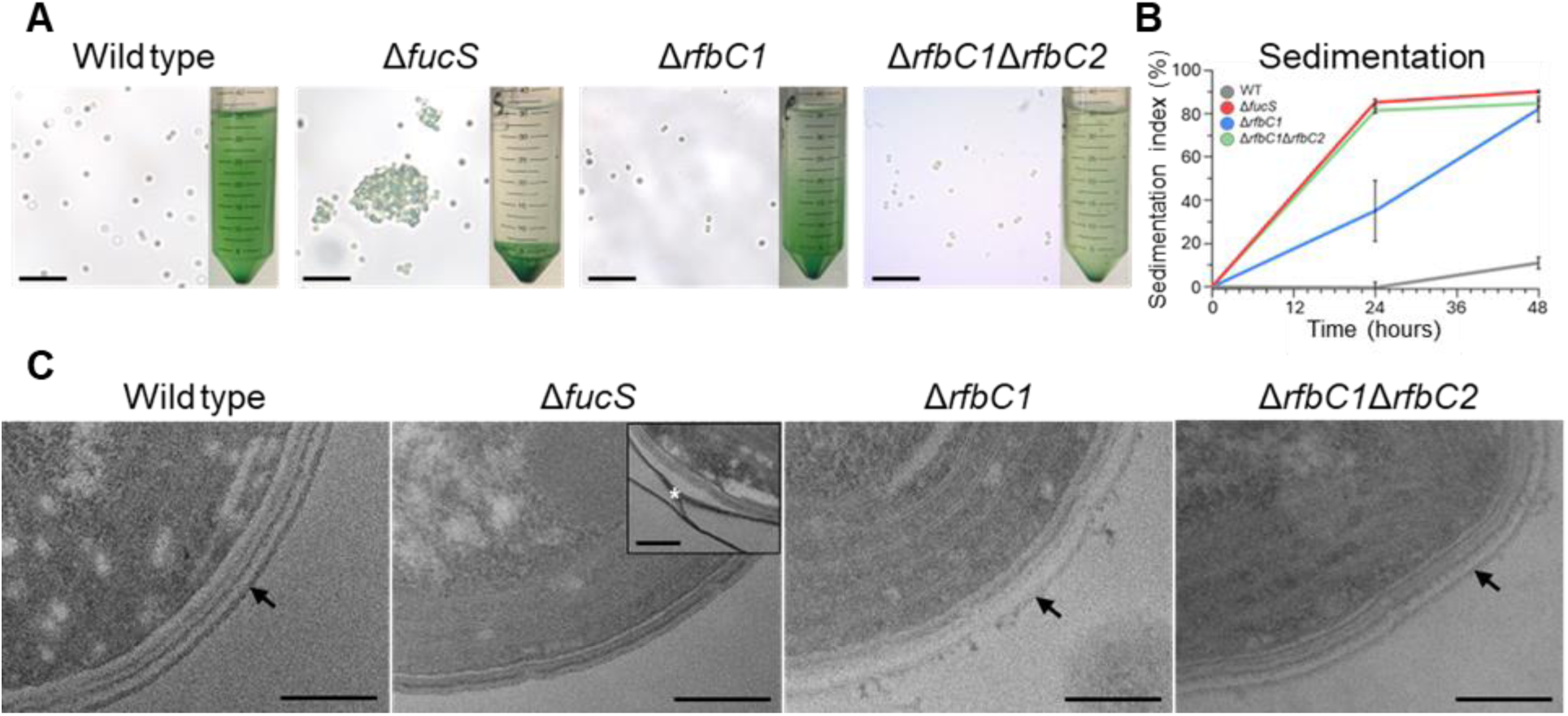
Aggregation/sedimentation and cell wall ultrastructure of *Synechocystis* sp. PCC 6803 wild type (WT), and Δ*fucS*, Δ*rfbC1* and Δ*rfbC1*Δ*rfbC2* strains. (A) Micrographs of the strains at low cell densities highlighting the clumping phenotype of Δ*fucS*, scale bar: 20 μm. Inserts highlight differences in sedimentation. (B) Sedimentation index (%) at 0, 24 and 48 h. Data represent means ± SD (*n* = 3). (C) Ultrastructure of the cell wall with the S-layer highlighted (arrow) and insert with the putative detached S-layer in Δ*fucS* also highlighted (asterisk). Scale bars: 200 nm. Cells were grown in BG11 medium at 30 °C under a 12 h light (25 μE m^−2^ s^−1^)/12 h dark regimen, with orbital shaking at 150 rpm.

Transmission electron microscopy confirmed the absence of the typical S-layer in the Δ*fucS* strain (Fig. 2C). However, in some cells, it was still possible to observe a string-like structure, with a similar electron density as the S-layer, suggesting that in Δ*fucS* the S-layer can be partially formed but is unable to remain attached to the cell wall (Fig. 2C, asterisk). Regarding Δ*rfbC1* and Δ*rfbC1*Δ*rfbC2*, both strains have a similar S-layer that seems to be slightly less electron dense and more uneven compared with the wild type.

Besides the obvious clumping phenotype and lack of S-layer, Δ*fucS* also exhibits a significant growth impairment compared with the wild type at 21 days: 14% and 23% regarding OD_730_ and chlorophyll *a*, respectively (Fig. 3A). The Δ*rfbC1* strain grows similarly to the wild type, while Δ*rfbC1*Δ*rfbC2* exhibits a significant growth impairment at 21 days: 13% in terms of OD_730_ and 32% regarding chlorophyll *a*. Concerning the total carbohydrate content neither Δ*fucS* nor Δ*rfbC1* showed differences compared with the wild type, while Δ*rfbC1*Δ*rfbC2* showed an increase of 35% (Fig. 3B). Regarding the relative amount of RPS, Δ*fucS* had no significant differences compared with the wild type, while Δ*rfbC1* and Δ*rfbC1*Δ*rfbC2* produced 35% and 27% less than the wild type, at the same time point (Fig. 3C). Additionally, both Δ*rfbC1* and Δ*rfbC1*Δ*rfbC2* produced 14% and 22% more CPS than the wild type, respectively (Fig. 3D).

**Fig. 3.**
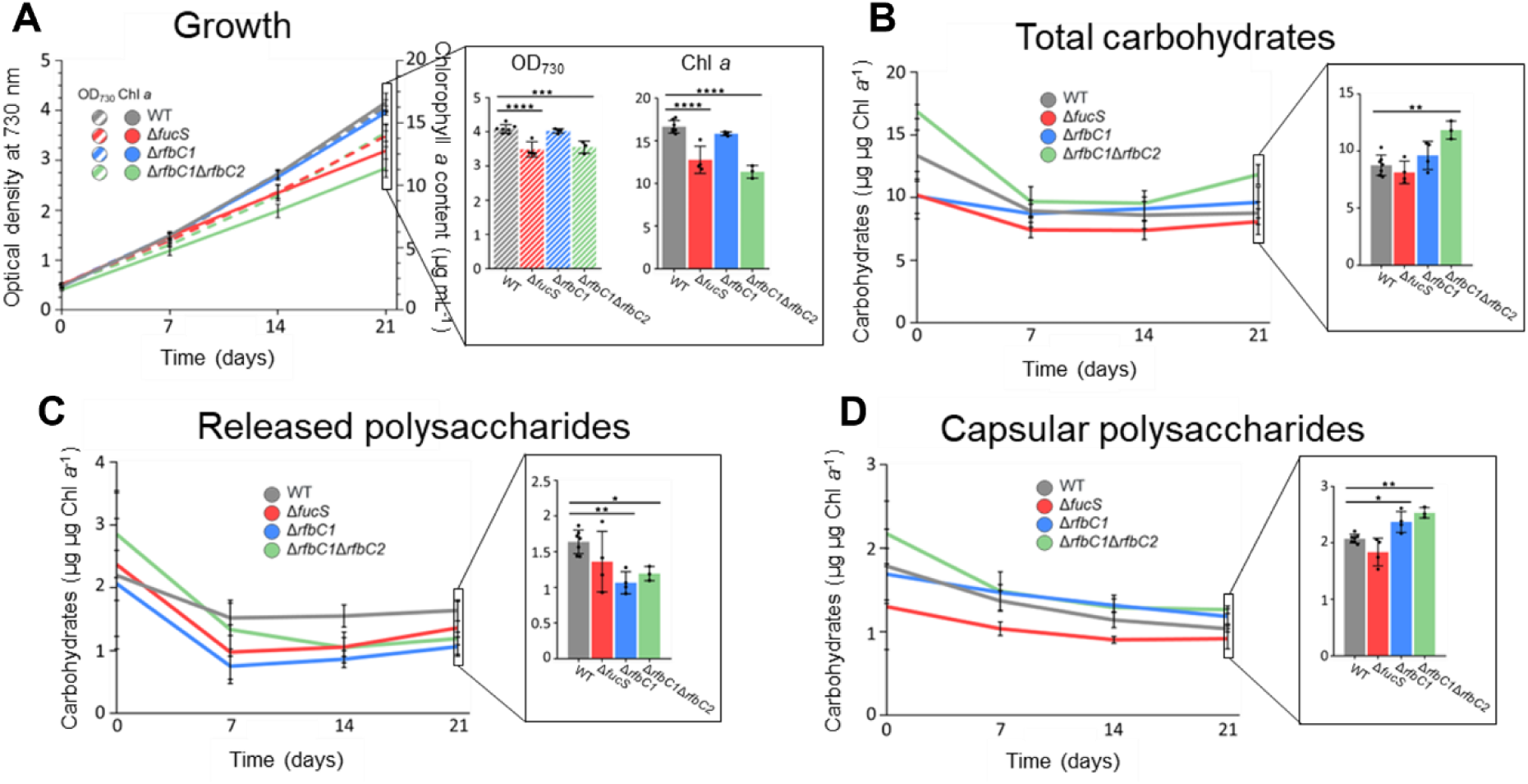
Growth, total carbohydrates, released polysaccharides and capsular polysaccharides of *Synechocystis* sp. PCC 6803 wild type (WT), and Δ*fucS*, Δ*rfbC1* and Δ*rfbC1*Δ*rfbC2* strains. (A) Growth (optical density at 730 nm) and μg of chlorophyll *a* per mL of culture, (B) production of total carbohydrates (C), released polysaccharides, and (D) capsular polysaccharides expressed as μg of carbohydrates per μg of chlorophyll *a*. Data represent means ± SD (*n* ≥ 3) and individual measurements are shown. Statistical analysis performed using one-way analysis of variance (ANOVA), followed by Dunnett’s multiple comparisons, is shown for the last time point. Significant differences are identified: *(*p* ≤ 0.05), **(*p* ≤ 0.01), ***(*p* ≤ 0.001), ****(*p* ≤ 0.0001).

### Monosaccharidic profiles of RPS and biomass

To evaluate if RPS composition also varies, the isolated polymers were hydrolysed and their monosaccharidic profile analysed. The Δ*fucS* RPS contain neither fucose nor rhamnose but have approximately 18% more hexoses, predominantly glucose. Another remarkable feature is the decrease in xylose: ∼53% less than the RPS from the wild type. Unexpectedly, the RPS from Δ*rfbC1* and Δ*rfbC1*Δ*rfbC2* still contain rhamnose, being both polymers quite similar to that of the wild type strain (Table 1).

**Table 1.** Monosaccharidic composition of the RPS from *Synechocystis* sp. PCC 6803 wild type, and Δ*fucS*, Δ*rfbC1* and Δ*rfbC1*Δ*rfbC2* strains expressed as molar %.

| Monosaccharides* | Wild type |  | *ΔfucS* |  | *ΔrfbC1* |  | *ΔrfbC1ΔrfbC2* |  |
| --- | --- | --- | --- | --- | --- | --- | --- | --- |
|  | % mol | SD | % mol | SD | % mol | SD | % mol | SD |
| Rha | 9.3 | 0.3 | nd. | 0.0 | 7.7 | 0.4 | 10.4 | 1.1 |
| Fuc | 10.6 | 0.5 | nd. | 0.0 | 9.5 | 0.5 | 11.7 | 1.8 |
| Rib | Traces | 0.1 | 1.1 | 0.1 | Traces | 0.1 | Traces | 0.1 |
| Ara | Traces | 0.1 | Traces | 0.1 | Traces | 0.1 | Traces | 0.1 |
| Xyl | 7.2 | 0.2 | **3.4** | 0.1 | 6.8 | 0.6 | 7.9 | 0.6 |
| Man | 9.4 | 1.0 | 7.6 | 0.4 | 9.3 | 0.4 | 8.6 | 1.1 |
| Gal | 3.1 | 0.2 | 6.6 | 0.7 | 4.5 | 1.6 | 3.1 | 0.1 |
| Glc | 52.6 | 1.5 | **68.9** | 0.5 | 53.3 | 0.8 | 47.7 | 2.1 |
| GalN | 1.8 | 0.0 | 1.9 | 0.1 | 2.7 | 0.0 | Traces | 0.0 |
| GlcN | 1.2 | 0.4 | 2.0 | 1.0 | 1.0 | 0.1 | 4.4 | 0.3 |
| Uronic Acids | 4.0 | 0.5 | 8.0 | 1.1 | 4.7 | 0.6 | 6.8 | 0.7 |

\*Rha: rhamnose; Fuc: fucose; Rib: ribose; Ara: arabinose; Xyl: xylose; Man: mannose; Gal: galactose; Glc: glucose; GalN: galactosamine; GlcN: glucosamine.
Traces < 1%; Not detected (nd.) = 0%. Sums may not be exactly 100% due to rounding.

To understand if rhamnose was present/absent in the CPS and/or lipopolysaccharides of the knockout strains and in particular in Δ*fucS*, a GFP-tagged protein that specifically binds to rhamnose was used to evaluate the amount of rhamnose attached to its cell wall relative to the wild type (for details see 2.11). The results obtained showed a striking ∼60% reduction in GFP signal for Δ*fucS* compared with the wild type, while Δ*rfbC1* showed a reduction of only ∼28% and for Δ*rfbC1*Δ*rfbC2* no statistical differences could be detected (Fig. 4).

**Fig. 4.**
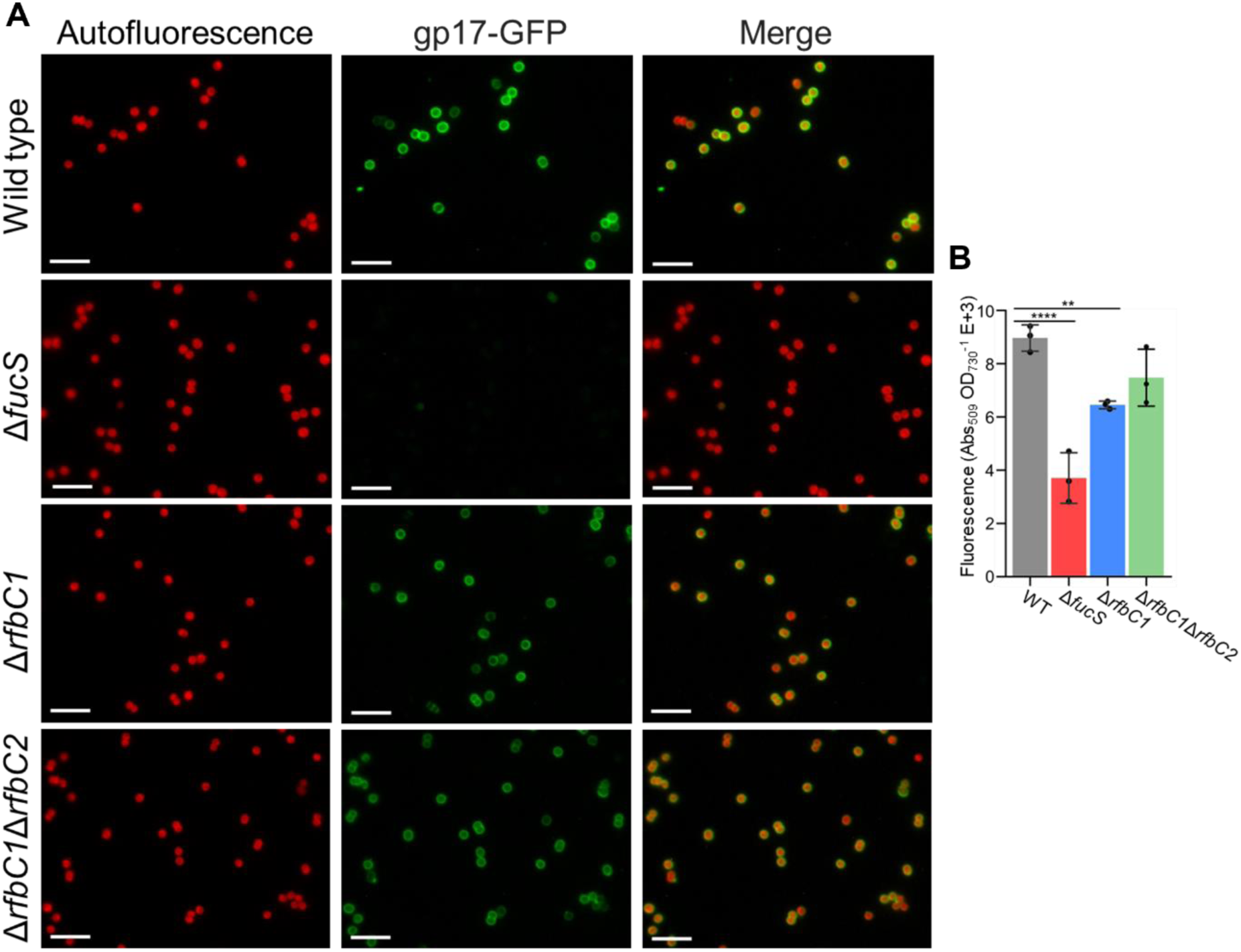
Detection of rhamnose on the cells surface of *Synechocystis* sp. PCC 6803 wild type (WT), and Δ*fucS*, Δ*rfbC1* and Δ*rfbC1*Δ*rfbC2* strains using a rhamnose binding protein tagged with GFP (gp17-GFP). (A) Micrographs depicting the red autofluorescence signal and the green signal from the gp17-GFP. The right panels show the merging of the other two micrographs. Scale bar: 10 μm. (B) Quantification of fluorescence associated with gp17-GFP bound to rhamnose at the surface of the cells. Data represents means ± SD (*n* = 3) and individual measurements are shown. Statistical analysis consists of one-way analysis of variance (ANOVA), followed by Dunnett’s multiple comparisons. Significant differences are identified: **(*p* ≤ 0.01) and ****(*p* ≤ 0.0001).

To determine if Δ*fucS* is only impaired in the incorporation of rhamnose into the EPS or also in the biosynthesis of this sugar, the monosaccharidic composition of the whole biomass was determined. The results obtained revealed no cytosolic fucose or rhamnose in the Δ*fucS* strain (Table 2). Since rhamnose was never detected by gas chromatography with a flame ionisation detector (GC-FID) in the Δ*fucS* strain, it seems reasonable to consider that the signal detected in Fig. 4B corresponds to background. The biomass sugars profile of Δ*rfbC1* is very similar to the wild type, while Δ*rfbC1*Δ*rfbC2* biomass shows a small decrease in glucose and an increase in all other sugars except the deoxyhexoses (Table 2). Once again, our results show no impairment in rhamnose biosynthesis for the two *rfbC* strains.

**Table 2.** Monosaccharidic composition of the biomass of *Synechocystis* sp. PCC 6803 wild type, and Δ*fucS*, Δ*rfbC1* and Δ*rfbC1*Δ*rfbC2* strains expressed as molar %.

| Monosaccharides* | Wild type | | $\Delta fucS$ | | $\Delta rfbC1$ | | $\Delta rfbC1\Delta rfbC2$ | |
| --- | --- | --- | --- | --- | --- | --- | --- | --- |
|  | % mol | SD | % mol | SD | % mol | SD | % mol | SD |
| Rha | 3.3 | 0.6 | nd. | 0.0 | 2.9 | 0.3 | 3.2 | 0.3 |
| Fuc | 2.5 | 0.9 | nd. | 0.0 | 2.7 | 0.1 | 2.6 | 0.4 |
| Rib | 4.4 | 0.7 | 5.8 | 0.8 | 4.6 | 0.5 | 7.8 | 0.7 |
| Ara | Traces | 0.2 | Traces | 0.0 | Traces | 0.1 | Traces | 0.1 |
| Xyl | 2.0 | 0.9 | Traces | 0.1 | 2.5 | 0.1 | 3.1 | 0.2 |
| Man | 3.0 | 0.6 | 3.6 | 0.2 | 3.9 | 0.6 | 6.1 | 2.6 |
| Gal | 7.7 | 1.6 | 13.9 | 2.0 | 12.2 | 0.4 | 11.7 | 1.4 |
| Glc | 68.8 | 2.5 | 65.4 | 3.6 | 61.4 | 1.9 | 51.4 | 3.5 |
| GalN | Traces | 0.1 | Traces | 0.1 | Traces | 0.0 | 1.5 | 1.4 |
| GlcN | 4.1 | 0.4 | 5.5 | 0.6 | 5.3 | 0.3 | 7.0 | 1.8 |
| Uronic Acids | 3.5 | 0.6 | 4.0 | 0.4 | 3.7 | 0.4 | 5.2 | 1.2 |

\*Rha: rhamnose; Fuc: fucose; Rib: ribose; Ara: arabinose; Xyl: xylose; Man: mannose; Gal: galactose; Glc: glucose; GalN: galactosamine; GlcN: glucosamine Traces < 1%; Not detected (nd.) = 0%. Sums may not be exactly 100% due to rounding.

### Transcriptional analysis of genes putatively involved in fucose and rhamnose biosynthesis and/or EPS production

To better understand the complex pathways involved in the synthesis of the deoxyhexoses, the transcript levels of the *slr1072* and *sll1212* (putative *gmd*, necessary for the biosynthesis of GDP-Fuc) and *sll1395* (putative *rfbD*, essential for the last step of dTDP-Rha biosynthesis), *sll1213* (*fucS*), *slr0985* (*rfbC1*) and *slr1933* (*rfbC2*) were analysed in the wild type, Δ*fucS,* Δ*rfbC1* and Δ*rfbC1*Δ*rfbC2* strains (Fig. 5 and S2). In addition, the transcript levels of other putative EPS-related genes present in the *rfbC1* operon were also analysed, namely *slr0982* (encoding a putative KpsT protein involved in EPS export), *slr0983* (encoding a putative RfbF transferase responsible for the activation of Glc-1-P to CDP-Glc in an alternative sugar pathway), and *slr1610* (encoding a hypothetical methyltransferase potentially involved in EPS methylation; Fisher et al., 2013).

**Fig. 5.**
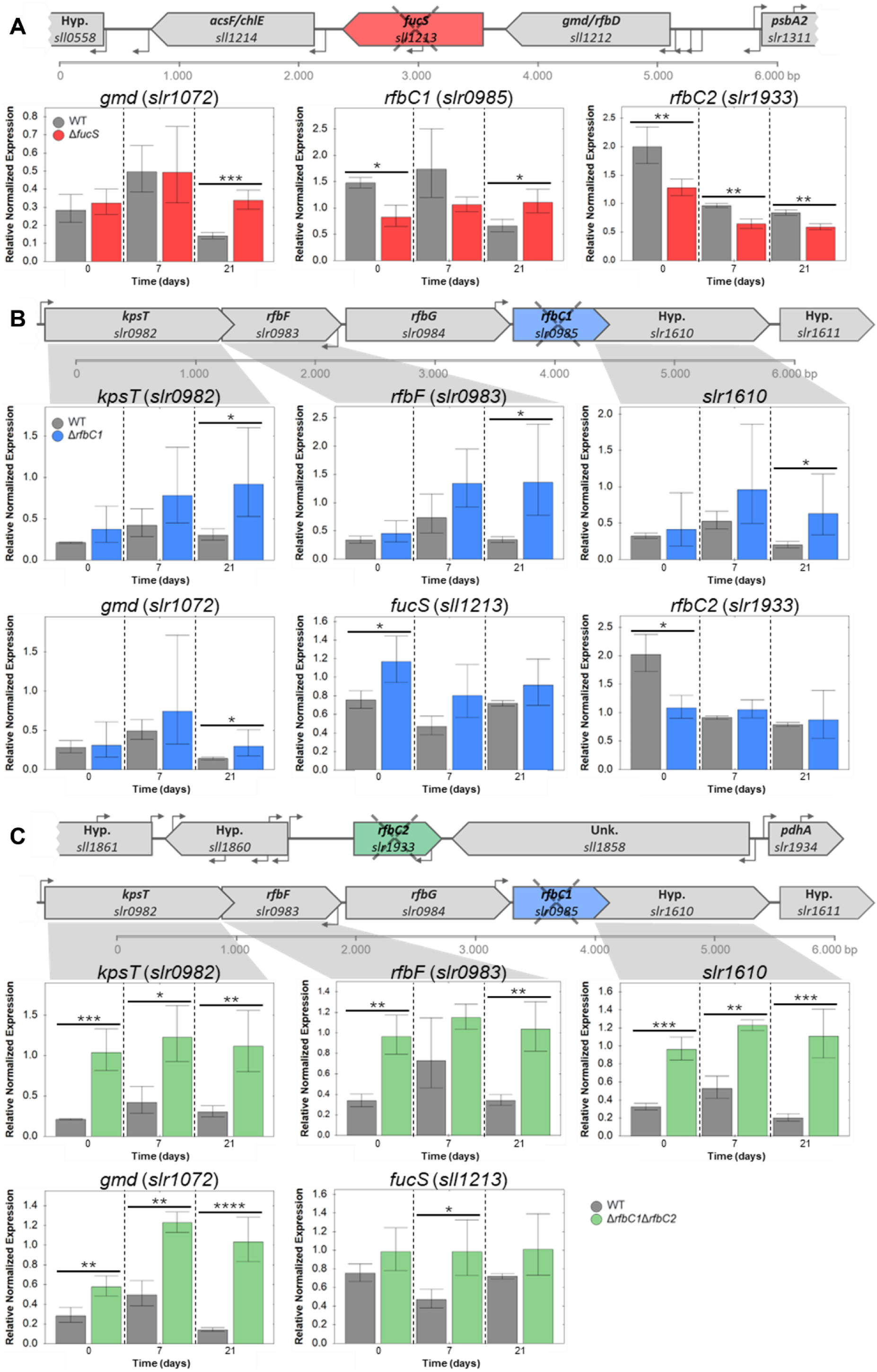
Analysis of the relative normalised expression of genes putatively related to the biosynthesis of deoxyhexoses and EPS in *Synechocystis* sp. PCC 6803 wild type (WT) and Δ*fucS*, Δ*rfbC1 and* Δ*rfbC1*Δ*rfbC2* strains. RNA was extracted from cells of the different strains collected at three time points: 0, 7 and 21 days. RT-qPCR analysis of (A) *gmd* (*slr1072*), *rfbC1* (*slr0985*) and *rfbC2* (*slr1933*) expression in *Synechocystis* sp. PCC 6803 wild type and Δ*fucS*; (B) *rfbB* (*slr0982*), *rfbF* (*slr0983*), *slr1610* (putative methyltransferase), *gmd* (*slr1072*), *fucS* (*sll1213*) and *slr1933* (*rfbC2*) expression in wild type and Δ*rfbC1*; and (C) *rfbB* (*slr0982*), *rfbF* (*slr0983*), *slr1610* (putative methyltransferase), *gmd* (*slr1072*) and *fucS* (*sll1213*) expression in wild type and Δ*rfbC1*Δ*rfbC2*. The normalised fold expression of the target genes relative to wild type is represented at each time point. Data from three biological and three technical replicates were normalised against two reference genes (*sll1212* and *sll1395*), and error bars represent the standard deviations. Statistical analysis was performed using *t*-test, and significant differences are identified: *(*p* ≤ 0.05), **(*p* ≤ 0.01), ***(*p* ≤ 0.001), ****(*p* ≤ 0.0001).

Among all the transcripts analysed, the levels of *sll1212* and *sll1395* (putative *gmd/rfbD* and *rfbD*, respectively) were stable in all strains and therefore used as reference genes (Fig. S2). In contrast, the results obtained showed that the transcript levels of the other putative *gmd* gene (*slr1072*) increased in all the knockout strains compared with the wild type. In Δ*fucS* and Δ*rfbC1* by 2.4- and 2.1-fold, respectively at 21 days (Fig. 5A and 5B), while in the double *rfbC* knockout strain the increase is more evident and over time (2.0-, 2.5- and 7.3-fold at 0, 7 and 21 days, respectively; Fig. 5C). In the Δ*rfbC1* and Δ*rfbC1*Δ*rfbC2* strains, all the analysed genes in the same operon as *rfbC1* have increased expression, more pronounced and at all time points in the double mutant. In Δ*rfbC1*, *kpsT* (*slr0982*), *rfbF* (*slr0983*) and *slr1610*, showed increased expression at 21 days of 3.0-, 4.0-, and 3.1-fold, respectively (Fig. 5B). In the Δ*rfbC1*Δ*rfbC2* strain, these three EPS-related targets were significantly upregulated with fold changes ranging from 2.3 to 5.5 (Fig. 5C). Furthermore, in the Δ*fucS* strain a decrease in the transcript levels of *rfbC2* was detected (Fig. 5A).

## DISCUSSION

In this work we aimed at better understand the inferred pathways for fucose and rhamnose biosynthesis in *Synechocystis* and how the deletion of genes encoding putative key enzymes impact EPS composition and production. For that, we targeted the epimerisation step of each deoxyhexose pathway. The GDP-Fuc pathway, and in particular the deletion of the *fucS* gene, had already been explored by others, who have described, among other things, impaired growth, cell aggregation, sedimentation, absence of the S-layer and EPS lacking fucose and rhamnose (Fisher et al., 2013; Matinha-Cardoso et al., 2022; Mohamed et al., 2005; Trautner and Vermaas, 2013). Our findings are in agreement with previous studies, but while the absence of the S-layer surrounding the cells was verified in the Δ*fucS* strain, sometimes it was possible to observe what appears to be an S-layer detaching from the cell surface (Fig. 2C). This observation, together with the results presented by Trautner and Vermaas (2013) that detected the S-layer protein (Sll1951) in Δ*fucS* culture supernatant, strongly suggests that this strain is capable of generating an S-layer but incapable of keeping it attached, most likely due to a fucose- or rhamnose-dependent anchor point. Regarding the other putative deoxyhexose pathway (dTDP-Rha), the deletion of *rfbC1* resulted in minor phenotypic alterations such as fast sedimentation rate and slightly altered S-layer, while the deletion of both putative copies (Δ*rfbC1*Δ*rfbC2*) had a stronger impact with an accentuated growth impairment and increase in sedimentation compared with both the wild type and the single deletion Δ*rfbC1* strains. More so, Δ*rfbC1*Δ*rfbC2* produced more total carbohydrates (per units of chlorophyll *a*). The *rfbC* deletions seem to affect, even if indirectly, the EPS assembly/export since both *rfbC* knockout strains have a shift in the production of EPS, with a decrease in RPS and an increase in CPS (Fig. 3C and 3D). It is reasonable to consider that this change is linked to the observed upregulation of the transcriptional unit of *rfbC1*, since all the targeted genes in the *rfbC1* operon become upregulated in the *rfbC* knockout strains (Fig. 5B and 5C). This could indicate that *rfbC1* and *rfbC2* gene products have redundant catalytic function or be similarly regulated. Among these upregulated genes are *kpsT* (*slr0982*), encoding an ATP-Binding Component of an ATP-Binding Cassette Transporter, and *slr1610*, encoding a methyltransferase, whose deletion has been shown to reduce EPS export (Fisher et al., 2013).

Strikingly, fucose and rhamnose are absent not only from the EPS of the Δ*fucS* strain, as previously shown (Fisher et al., 2013), but also from its biomass indicating that this strain is indeed incapable of synthesising rhamnose. It is intriguing, however, how the deletion of *fucS* leads to the inability of *Synechocystis* to produce rhamnose. To the best of our knowledge, there is no naturally occurring system for rhamnose biosynthesis that has fucose as precursor, raising the question of how these two sugar pathways are linked. Many monosaccharide pathways, and in particular the respective nucleosiltransferases, have been shown to be under the regulation of a myriad of molecules, such as end product sugars of the pathway, other sugars, and other nucleoside-derived compounds (Alphey et al., 2013; Blankenfeldt et al., 2000; Hulen, 2023; Zheng et al., 2023). An example of this is RfbA, the nucleosiltransferase of the dTDP-Rha pathway, which in several organisms has been shown to be allosterically regulated by a variety of mechanisms, such as analogue sugars and compounds derived from dTDP (Alphey et al., 2013; Zheng et al., 2023). With this in mind, one can hypothesise that the absence of *fucS* leads to an accumulation of GDP-4-keto-6-deoxy-D-mannose (not too dissimilar to dTDP-4-keto-6-deoxy-L-mannose, precursor of dTDP-Rha), which in turn could interfere with the catalytic function of RfbA culminating in the absence of rhamnose. However, it is important to notice that GDP-D-rhamnose (GDP-Rha) has been shown not to inhibit RfbA in other bacteria, such as *Escherichia coli*, *Pseudomonas aeruginosa*, *Salmonella enterica* and *Mycobacterium tuberculosis* (Zheng et al., 2023).

Additionally, and contrary to what was expected, rhamnose residues are present in both RPS and biomass of the Δ*rfbC1* and Δ*rfbC1*Δ*rfbC2* strains at similar amounts to the ones observed for the wild type. Under the assumption that *Synechocystis* produces dTDP-Rha, these results indicate that there must be another *rfbC* gene yet to be identified. However, in some rare cases, GDP-4-keto-6-deoxy-D-mannose is also a precursor of GDP-Rha as for example in *Xanthomonas campestris* and *Pseudomonas aeruginosa* (Mäki et al., 2002; Vorhölter et al., 2001). It has also been previously described that in *Synechocystis* the sequence of *slr0583* is similar to that of *fucS*, and while the biological function of the corresponding protein remains unknown, Mohamed et al. (2005) noted that *slr0583* is also similar to a *rmd* gene in *Xanthomonas campestris* that is essential for the biosynthesis of GDP-Rha. Therefore, one cannot discard the possibility that *Synechocystis* could produce GPD-Rha *via* a pathway closely related to the pathway of GDP-Fuc.

Interestingly, the transcript levels of *sll1212* (the putative *gmd/rfbD* immediately upstream of *fucS*; Fig. 1B) did not vary significantly in all strains, although this gene could be considered the prime *gmd* candidate as observed in many other organisms (Mäki and Renkonen, 2004). In contrast, the transcript levels of the other *gmd* candidate (*slr1072*) were upregulated in all strains, suggesting that *slr1072* is important for the biosynthesis of GDP-Fuc, dTDP-Rha or other closely related pathway such as GDP-Rha. If *slr1072* does encode a Gmd, its striking upregulation in Δ*rfbC1*Δ*rfbC2* could be explained by an activation of a GDP-Rha pathway due to the inability to produce dTDP-Rha.

In conclusion, deoxyhexoses biosynthetic pathways in *Synechocystis* are more complex, and/or have more players, than initially anticipated. Here we demonstrated that *fucS* (*sll1213*) deletion impairs both fucose and rhamnose biosynthesis and that the putative *rfbC* (*slr0985* and *slr1933*) deletion(s) do not impair rhamnose biosynthesis despite affecting EPS export. Additionally, increased expression of the putative *gmd* (*slr1072*) in Δ*fucS* and Δ*rfbC* strains suggests that its gene product might have a important role in the deoxyhexoses’ pathways. More comprehensive studies are necessary to identify the key players as well as their regulatory mechanisms.

## MATERIALS AND METHODS

### Bacterial strains and standard growth conditions

The cyanobacterium *Synechocystis* sp. PCC 6803 wild type (sub-strain GT-Kazusa; glucose tolerant, with S-layer and non-motile) and strains were cultivated in BG11 medium (Stanier et al., 1971) at 30 °C under a 12 h light (25 μmol photons m^−2^ s^−1^)/12 h dark regimen with orbital shaking (150 rpm). For solid medium, BG11 medium was supplemented with 1.5% (w/v) Noble agar (Difco™), 0.3% (w/v) sodium thiosulfate, and 10 mM TES [N-tris(hydroxymethyl)methyl-2-aminoethanesulfonic acid]-potassium hydroxide (KOH) buffer (pH 8.2). For the selection and maintenance of strains, BG11 medium was supplemented with kanamycin (Km; Merck Millipore) up to 400 μg mL^−1^ and/or chloramphenicol (Cm; Merck Millipore) up to 75 μg mL^−1^. The *Escherichia coli* TOP10 (Thermo Fisher Scientific) was cultivated at 37 °C in LB medium (Bertani, 1951) supplemented with ampicillin (100 μg mL^−1^; Amp; Merck Millipore), Km (50 μg mL^−1^), or Cm (25 μg mL^−1^).

### Cyanobacterial DNA extraction and recovery

Cyanobacterial genomic DNA was extracted by the phenol-chloroform method described previously (Tamagnini et al., 1997). Agarose gel electrophoresis was performed according to standard protocols (Sambrook and Russel, 2001), and the DNA fragments were isolated from gels using the Monarch^®^ DNA gel extraction kit (New England Biolabs), and from enzymatic assay mixtures or PCR mixtures using the Monarch^®^ PCR & DNA cleanup kit (New England Biolabs).

### Plasmid construction for *Synechocystis* transformation

The *Synechocystis* sp. PCC 6803 chromosomal regions flanking *rfbC1* (*slr0985*) and *rfbC2* (*slr1933*) were amplified by PCR using specific oligonucleotide primers (Table S1). An overlapping region containing a *Stu*I restriction site was included in primers slr0985.5I and slr0985.3I and a *Xma*I restriction site was included in primers slr1933.5R and slr1933.3F for cloning purposes. For each gene, the purified PCR fragments were fused by “overlap PCR”. The resulting products were purified and cloned into the vector pGEM^®^-T Easy (Promega), creating plasmids pGDslr0985 and pGDslr1933. A selection cassette containing the *nptII* gene (resistance to neomycin and kanamycin) was excised from plasmid pKm.1 (Pinto et al., 2015) with the restriction enzyme *Sma*I or *Xma*I (Thermo Fisher Scientific) and *cat* (resistance to chloramphenicol) was amplified from pSEVA351 (Silva-Rocha et al., 2013) by PCR using primers containing a *Xma*I restriction site. Subsequently, the selection cassettes were cloned into the *Xma*I restriction site of the plasmids to form pGDslr0985.Km and pGDslr1933.Cm.

### Generation of Synechocystis ΔrfbC1 and ΔrfbC1ΔrfbC2

Δ*fucS* (Δ*sll1213*) was kindly provided by P. Oliveira (Matinha-Cardoso et al., 2022). To generate the other strains, *Synechocystis* sp. PCC 6803 cultures were grown until the optical density at 730 nm (OD_730_) reached ∼0.8, and cells were harvested by centrifugation and suspended in BG11 medium to an OD_730_ of ∼2.5. Five hundred microliters of cells were incubated with 20 μg mL^−1^ plasmid DNA for 5 h before spreading them onto Immobilon-NC membranes (0.45 μm pore size; Merck Millipore) resting on solid BG11 plates, and kept at 26 °C under 16 h light/8 h dark for 24 h. Then, membranes were transferred to selective plates containing 10 μg mL^−1^ Km or 12.5 μg mL^−1^ Cm. After one week, transformants were transferred to plates containing 20 μg mL^−1^ Km or 25 μg mL^−1^ Cm. For complete segregation, colonies were grown in increasing antibiotic concentrations.

### Strain confirmation by PCR and Southern Blot

Segregation was confirmed by PCR, using the primer pairs listed in Table S1 and depicted in Fig. S1. Each PCR reaction contained 2-10 ng of genomic DNA, 0.25 µM of each primer, 200 µM of dNTPs mix (Promega), 1x Green GoTaq^®^ Flexi buffer, 1.5 mM of MgCl_2_ and 0.5 U of GoTaq^®^ G2 Flexi DNA Polymerase (Promega). For the PCRs an initial denaturation of 95 °C for 5 min was performed, as well as a final extension at 72 °C for 7 min. Strain’s segregation was further confirmed by Southern blot. To confirm deletion of *rfbC1* (*slr0985*) approximately 2-3 μg of genomic DNA of *Synechocystis* sp. PCC 6803 wild type, Δ*rfbC1* and Δ*rfbC1*Δ*rfbC2* was digested with *Dra*I (Thermo Fisher Scientific). For the deletion of *rfbC2* (*slr1933*), 2-3 μg of genomic DNA of wild type and Δ*rfbC1*Δ*rfbC2* was digested with *Nco*I (Thermo Fisher Scientific). The DNA fragments were separated on a 0.8% (w/v) agarose gel and transferred by vacuum (5 inHg, Bio-Rad, Vacuum regulator) onto an Amersham™ Hybond™-N+ (Cytiva) with a Model 785 Vacuum Blotter (Bio-Rad). For the transfer, the DNA was depurinated and denaturated with 0.25 M HCl (20 min) followed by 0.5 M NaOH 1.5 M NaCl (20 min). The gel was soaked in neutralising buffer (1 M Tris-HCl pH 7.5, 1.5 M NaCl) for 20 min and subsequently in 20x SSC (3 M NaCl, 0.3 M sodium citrate pH 7.0) for 30 min before being equilibrated in 6x SSC for 5 min. The membrane was removed from the Vacuum Blotter equilibrated in 6x SSC for 5 min and cross-linked at 80 mJ cm^-2^. For hybridisation, probes covering the flaking regions 5’ of *rfbC1* and 3’ of *rfbC2* were amplified by PCR using primer pairs slr0985_SB_Fwd and slr0985_SB_Rev, and slr1933.3F and slr1933.3R (respectively, Table S1). Probes were then labelled using the DIG DNA labelling kit (Roche), according to the manufacturer’s instructions. Hybridisations were performed overnight at 60 °C in a solution containing 2% (w/v) Blocking Reagent (Roche), 0.1% (w/v) N-lauroylsarcosine, 3x SSC and 0.02% (w/v) SDS. Subsequently, the membrane was immersed in Low Stringency Buffer [2x SSC, 0.1% (w/v) SDS] at room temperature for 5 min (twice) and in High Stringency Buffer [0.5x SSC, 0.1% (w/v) SDS] at 60 °C for 15 min (twice). The membrane was placed in Washing Buffer [0.1 M Maleic Acid Buffer, pH 7.5 with 0.3% (v/v) Tween-20] for 2 min and then incubated in Blocking Reagent 1% for 30 min followed by Blocking Reagent 1% (w/v) with Anti-DIG antibody at 1:10000 for 30 min before being washed with 2x Washing Buffer for 15 min. Finally, the membrane was incubated with Detection Buffer (0.1 M Tris-HCl, 0.1 M NaCl, pH 9.5) for 5 min followed by incubation with CDP-Star (Roche) for 5 min and revealed using a ChemiDoc Imager (Bio-Rad).

### Growth assessment

*Synechocystis* cultures were inoculated to an OD_730_ of 0.5 in 150 mL of BG11 medium in 250 mL Erlenmeyer flasks and grown for 21 days, under standard growth conditions. Growth measurements were performed every 7 days by monitoring the OD_730_ (Anderson and McIntosh, 1991) using a spectrophotometer (Shimadzu Uvmini-1240, Shimadzu Corporation) and determining the chlorophyll *a* content as described previously (Meeks and Castenholz, 1971). All experiments were performed with three technical and three biological replicates.

### Determination of total carbohydrate, RPS, and CPS contents

Total carbohydrates and RPS contents were determined as described previously by Mota et al. (2013) and CPS as described by Santos et al. (2021) using the phenol-sulfuric acid method (DuBois et al., 1956). All contents were expressed as μg of carbohydrates per μg of chlorophyll *a*. All experiments were performed with three technical and three biological replicates.

### Sedimentation index

Cell culture sedimentation was quantified by measuring the OD_730_ of the cultures at 0 h, 24 h, and 48 h. For that, 4 mL of homogeneous culture at OD_730_ of 0.8 was transferred to plastic cuvettes (Fisher Scientific) and left to sediment for 48 h at room temperature. Sedimentation index was calculated using the equation [(OD_730_i - OD_730_t)x(OD_730_i)^-1^]x100 where OD_730_i is the measurement of the initial OD_730_ (at 0 h) and OD_730_t is the OD_730_ at each other time point.

### Optical microscopy

Cells were observed directly using an Axio Lab.A1 light microscope and micrographs were acquired with an Axiocam Erc 5 s camera using the ZEN 2.6 software (ZEISS).

### Transmission electron microscopy

Cells were fixed directly in culture medium with final concentrations of 2.5% (v/v) glutaraldehyde and 2% (v/v) paraformaldehyde in 0.05 M sodium cacodylate buffer (pH 7.2) (overnight), washed three times in double-strength sodium cacodylate buffer followed by post-fixation with 2% (v/v) osmium tetroxide and 1.5% (w/v) potassium ferrocyanide in 0.1 M sodium cacodylate buffer (pH 7.4) for 3 h and washed three times with MilliQ^®^ grade water. Samples were then suspended and incubated in 2% (v/v) tannic acid for 1 h and washed three times with MilliQ^®^ grade water. Pellets were suspended in 2% (v/v) uranyl acetate and left to incubate overnight at 4 °C and washed three times in MilliQ^®^ grade water. Warm HistoGel™ (Fisher Scientific) was added to the cell pellets and left to cool at 4 °C until solid. Sample was gradually dehydrated with solutions of increasing concentration of ethanol in MilliQ^®^ grade water and a final step with propylene oxide. Embedding was performed gradually with increasing concentration of EPON resin in propylene oxide until pure resin was added and left to incubate for 48 h at 60 °C until completely solid. Ultrathin sections were then mounted on 300 mesh copper grids and samples were stained with 2% (v/v) uranyl acetate and 2% (w/v) lead citrate [Reynolds’ method (Reynolds, 1963)] for 15 min each. Samples were examined under a JEM-2100-HT TEM (JEOL Ltd.) operating at 80 kV and equipped with a fast-readout “OneView” 4k × 4k CCD camera that operates at 25 fps (300 fps with 512 × 512 pixel).

### Rhamnose binding assay

Cell surface rhamnose was quantified using a bacteriophage receptor binding protein (gp17) fused to GFP (gp17-GFP) provided by M. Loessner (Bielmann et al., 2015). Cultures were inoculated in 10 mL BG11 medium at OD_730_ of 0.2 and incubated at 30 °C under a 12 h light (25 μmol photons m^−2^ s^−1^)/12 h dark regimen with orbital shaking (150 rpm) until an OD_730_ = 1.5. Cultures were centrifuged for 10 min at 4470 *g* and cell pellets suspended in fresh BG11 medium to a final concentration of OD_730_ = 5. One hundred µL of culture was transferred to a 96-well microplate (Greiner Bio-One) and cells washed with phosphate buffered saline (PBS) (10 min at 3200 *g*). Then, cells were incubated in PBS complemented with gp17-GFP at 17 µg mL^−1^ for 20 min in the dark. After that, cells were washed 3 times with PBS to remove unbound protein. To prevent cell clumping, cells were resuspended in 20 mM Tris-HCl (pH 7.8) with 4% (w/v) SDS. OD_730_ and GFP fluorescence were measured using a CLARIOstar Plus plate reader (BMG Labtech). Fluorescence data was normalised by OD_730_. All experiments were performed with three technical and three biological replicates. Images were collected with an Olympus BX53 (Olympus Corporation) fluorescence microscope with ISO set to 400. Autofluorescence micrographs were captured using the filter for tetramethylrhodamine (TRITC) and an exposure of 714.2 ms, while the micrographs of the gp17-GFP fluorescence were obtained using filter for fluorescein (FITC) and an exposure of 957.4 ms. Images were then processed using Fiji™ (version 2.15.0, Schindelin et al., 2012).

### Neutral sugars and uronic acids analysis

To determine the monosaccharidic composition of the RPS, cultures at OD_730_ = 3 were processed as described by Flores and Tamagnini (2019). To determine the monosaccharidic composition of the biomass, cell cultures at OD_730_ = 1.5 were centrifuged and washed with BG11. Cell pellets were left to dry at 55 °C for 48 h. Both freeze-dried RPS and biomass (1 to 2 mg) were processed and analysed as detailed by Aveiro et al. (2020). In short, all samples were subjected to a prehydrolysis with 0.2 mL of 72% (v/v) H_2_SO_4_ for 3 h at room temperature, followed by 2.5 h hydrolysis with 1 M H_2_SO_4_ at 100 °C. Neutral sugars were determined by converting the hydrolysed sugars in alditol acetates, using 2-desoxyglucose as internal standard. Then, gas chromatography with a flame ionisation detector (GC-FID; Perkin-Elmer Clarus 400) equipped with a DB-225 capillary column (Agilent J&W GC columns) was used to identify the alditol acetates. The uronic acids content was determined by the m-phenylphenol colorimetric method, and d-Galacturonic acid solutions (0–80 µg mL^−1^) were used to construct a calibration curve. The analyses were performed in triplicate.

### RNA extraction, cDNA synthesis and transcription analysis by RT-qPCR

For RNA extraction, cultures were inoculated to OD_730_ = 0.5 in a final volume of 150 mL of BG11 medium in 250 mL Erlenmeyer flasks and grown for 21 days. Samples were collected at 0, 7 and 21 days and cell pellets were treated as previously detailed by Ferreira et al. (2018). RNA concentration, purity and integrity was checked as stated by Ferreira et al. (2018) and for cDNA synthesis, 1 µg of total RNA was transcribed with the iScript^TM^ Reverse Transcription Supermix for RT-qPCR (Bio-Rad) in a final volume of 20 µL, following the manufacturer’s instructions. Five-fold standard dilutions of the cDNAs were made (1/5, 1/25, 1/125 and 1/625) and stored at -20 °C.

RT-qPCRs were performed on Hard-Shell 384-Well PCR Plates (thin wall, skirted, clear/white) covered with Microseal^®^ B adhesive seal (Bio-Rad). The reactions (10 µL) were manually assembled and contained 0.125 μM of each primer (Table S2), 5 μL of iTaq™ Universal SYBR^®^ Green Supermix (Bio-Rad) and 1 μL of template cDNA (dilution 1/5). The PCR protocol used was: 3 min at 95 °C followed by 40 cycles of 30 s at 95 °C, 30 s at 56 °C and 30 s at 72 °C. In the end, a melting curve analysis of the amplicons (10 s cycles between 65 – 95 °C with a 0.5 °C increment per cycle) was carried out. Standard dilutions of the cDNA were used to check the relative efficiency and quality of primers, and negative controls (no cDNA template) were included. RT-qPCRs were performed with three biological and three technical replicates of each cDNA sample in the CFX384 Touch™ Real-Time PCR Detection System (Bio-Rad). The data obtained were analysed using the Bio-Rad CFX Maestro™ 1.1 software (Bio-Rad), implementing an efficiency-corrected delta-delta Cq method (ΔΔCq). The genes *sll1395* and *sll1212* were validated as reference genes for data normalisation using the reference gene selection tool available in the Bio-Rad CFX Maestro™ software (version 2.3, Bio-Rad). Standard *t*-test was performed using the same software, and tests were considered significant if *p* < 0.05.

### Statistical analysis

Data were statistically analysed with GraphPad Prism (version 8.0.1, GraphPad Software) using analysis of variance (ANOVA), followed by Dunnett’s multiple comparisons test.

## ACKNOWLEDGEMENTS

This work was conducted within the framework of the scholarship 2020.08663.BD (J.P.), funded by National Funds through FCT – Fundação para a Ciência e a Tecnologia, I.P., and by ESF – European Social Fund through the POCH – Programa Operacional Capital Humano within the framework of PORTUGAL2020, namely through the NORTE 2020 – Programa Operacional Regional do Norte. It was also conducted withing the context of the Molecular and Cell Biology doctoral program from ICBAS. FCT – Fundação para a Ciência e a Tecnologia, I.P. also funded this work under the project UIDB/04293/2020. FCT is also acknowledged for the Assistant Researcher contracts CEECIND/00259/2017 (C.C.P.) and 2020.01953.CEECIND (F.P.). This work was also in part developed within the scope of the project CICECO-Aveiro Institute of Materials (UIDB/50011/2020, UIDP/50011/2020 & LA/P/0006/2020) and LAQV-REQUIMTE (UIDB/50006/2020, UIDP/50006/2020), financed by national funds through the FCT/MEC (PIDDAC).

We wish to express our gratitude to Dr. Paulo Oliveira and Jorge Matinha-Cardoso for the Δ*fucS* strain used in this study and, in particular, to Dr. Paulo Oliveira for his valuable inputs throughout this work. We would also like to thank the i3S Scientific Platforms: Biochemical and Biophysical Technologies, Genomics, and Cell Culture and Genotyping. Especially Cell Culture and Genotyping for their support in the RT-qPCR experiments. Additionally, we thank Dr. Ana Rita Malheiro for her help regarding the transmission electron microscopy (INL Electron Microscopy and X-Rays Facility).

